# The major subunit of widespread competence pili exhibits a novel and conserved type IV pilin fold

**DOI:** 10.1101/2020.02.12.945410

**Authors:** Devon Sheppard, Jamie-Lee Berry, Rémi Denise, Eduardo P. C. Rocha, Steve Matthews, Vladimir Pelicic

**Author notes:** To whom correspondence should be addressed: Vladimir Pelicic: Imperial College London, Medical Research Council Centre for Molecular Bacteriology and Infection, London SW7 2AZ, United Kingdom.

## Abstract

Type IV filaments (T4F), which are helical assemblies of type IV pilins, constitute a superfamily of filamentous nanomachines virtually ubiquitous in prokaryotes that mediate a wide variety of functions. The competence (Com) pilus is a widespread T4F, mediating DNA uptake (the first step in natural transformation) in bacteria with one membrane (monoderms), an important mechanism of horizontal gene transfer. Here, we report the results of genomic, phylogenetic, and structural analyses of ComGC, the major pilin subunit of Com pili. By performing a global comparative analysis, we show that Com pili genes are virtually ubiquitous in Bacilli, a major monoderm class of Firmicutes. This also revealed that ComGC displays extensive sequence conservation, defining a monophyletic group among type IV pilins. We further report ComGC solution structures from two naturally competent human pathogens, *Streptococcus sanguinis* (ComGC_SS_) and *Streptococcus pneumoniae* (ComGC_SP_), revealing that this pilin displays extensive structural conservation. Strikingly, ComGC_SS_ and ComGC_SP_ exhibit a novel type IV pilin fold that is purely helical. Results from homology modelling analyses suggest that ComGC unusual structure is compatible with helical filament assembly. Because ComGC displays such a widespread distribution, these results have implications for hundreds of monoderm species.

## Introduction

Filamentous nanomachines composed of type IV pilins are virtually ubiquitous in Bacteria and Archaea (1), to which they confer a variety of unrelated functions including adhesion, motility, protein secretion, DNA uptake. These type IV filaments (T4F) are assembled by conserved multi-protein machineries, which further underlines their phylogenetic relationship (2).

Much of our current understanding of this superfamily of nanomachines comes from the study of type IV pili (T4P), the best characterised T4F (1). In brief, T4P are μm-long and thin surface-exposed filaments, which are polymers of type IV pilins. Type IV pilins (simply named pilins hereafter) are defined by an N-terminal sequence motif known as class III signal peptide (3). This motif – IPR012902 entry in the InterPro database (4) – consists of a hydrophilic leader peptide ending with a tiny residue (Gly or Ala), followed by a tract of 21 mostly hydrophobic residues, except for a negatively charged Glu_5_. This hydrophobic tract represents the N-terminal portion (α1N) of an extended α-helix of ~50 residues (α1), which is the universally conserved structural feature of type IV pilins (3). Although some small pilins consist solely of this extended α-helix (5), most pilins have a globular head consisting of the C-terminal half of α1 (α1C) packed against a β-sheet composed of several antiparallel β-strands, which gives them their typical “lollipop” 3D architecture (3). Upon translocation of prepilins across the cytoplasmic membrane (CM), where they remain embedded via their protruding hydrophobic α1N, the leader peptide is processed by an integral membrane aspartic acid protease named prepilin peptidase (IPR000045) (6). Processing primes pilins for polymerisation into filaments. Filament assembly, which remains incompletely understood, is mediated by a multi-protein machinery in the CM, centred on an integral membrane platform protein (IPR003004) and a cytoplasmic extension ATPase (IPR007831) (1). As revealed by recent cryo-EM structures of several T4P (7,8), filaments are right-handed helical polymers where pilins are held together by extensive interactions between their α1 helices, which are partially melted and run approximately parallel to each other within the filament core.

One of the key functional roles of T4F is their involvement in natural transformation in prokaryotes, the ability of species defined as “competent” to take up exogenous DNA across their membrane(s) and incorporate it stably into their genomes (9). This widespread property in bacteria (10) is key for horizontal gene transfer, an important factor in bacterial evolution and the spread of antibiotic resistance. T4F are involved in the very first step of natural transformation, *i.e.* binding of free extracellular DNA and its translocation close to the CM (9). DNA is subsequently bound by the DNA receptor ComEA and further translocated across the CM through the ComEC channel (9). In diderm competent species, the T4F involved in DNA uptake is a subtype of T4P, known as T4aP (11), which rapid depolymerisation is powered by the retraction ATPase PilT (IPR006321), generating exceptionally large tensile forces (12). In brief, T4aP bind DNA directly, via one of their major or minor (low abundance) pilin subunits (13), and then are retracted by PilT, bringing DNA to the ComEA receptor (14). In monoderm competent species, DNA uptake is mediated by a distinct T4F named competence (Com) pilus (9), much less well characterised than T4P. Com pili are composed mainly of the major pilin (ComGC) (15,16), and are assembled by a simple machinery composed of four minor pilins (ComGD, ComGE, ComGF, ComGG), a prepilin peptidase (ComC), an extension ATPase (ComGA) and a platform protein (ComGB) (17,18). Filaments morphologically similar to T4aP, several μm in length and 60 Å in width, have been observed in *S. pneumoniae* (15,19).

How Com pili are assembled, bind DNA and presumably retract in the absence of a PilT retraction motor is not understood. One important limitation is the absence of high-resolution structural information. Therefore, in the present study, we have focused on ComGC, the major subunit of the Com pilus. We report (i) a global comparative and phylogenetic analysis of ComGC, and (ii) 3D structures for two orthologs, ComGC_SP_ from the model competent species *S. pneumoniae* and ComGC_SS_ from *S. sanguinis*, a common cause of infective endocarditis in humans that has recently emerged as a monoderm model for the study of T4F. Finally, we discuss the general implications of these findings.

## Results

### Com pili genes are almost ubiquitous in monoderm Bacilli, including the T4F model S. sanguinis

So far, Com pili have been mainly studied in two model competent species: *B. subtilis* and *S. pneumoniae*. *S. sanguinis* is a naturally competent species that has recently emerged as a monoderm T4F model since it expresses retractable T4aP (20). Functional analysis of *S. sanguinis* T4aP showed that they are dispensable for DNA uptake, which is instead mediated by Com pili since competence was abolished in a *∆comGB* mutant (21). A closer inspection of *S. sanguinis* genome revealed that all the genes encoding the Com pilus are present. These genes are organised in two loci (Fig. 1A), *comC* and the *comGABCDEDFG* operon, showing perfect synteny with the corresponding loci in model competent species (22,23). Multiple sequence alignments of the corresponding proteins with orthologs in *B. subtilis* and *S. pneumoniae* showed extensive conservation (Table S1). Detailed sequence analysis of the N-termini of the five ComG pilins identified clear class III signal peptides (Fig. 1B), *i.e.* short (8-15 residues) and hydrophilic leader peptides ending with an Ala, followed by a tract of 21 mostly hydrophobic residues. ComGG is the only pilin that does not have a negatively charged Glu5 and displays a non-canonical class III signal peptide (Fig. 1B), which is not identified by InterPro or PilFind that is dedicated to the prediction of type IV pilins (24). This is a conserved property for ComGG orthologs.

**Fig 1.**
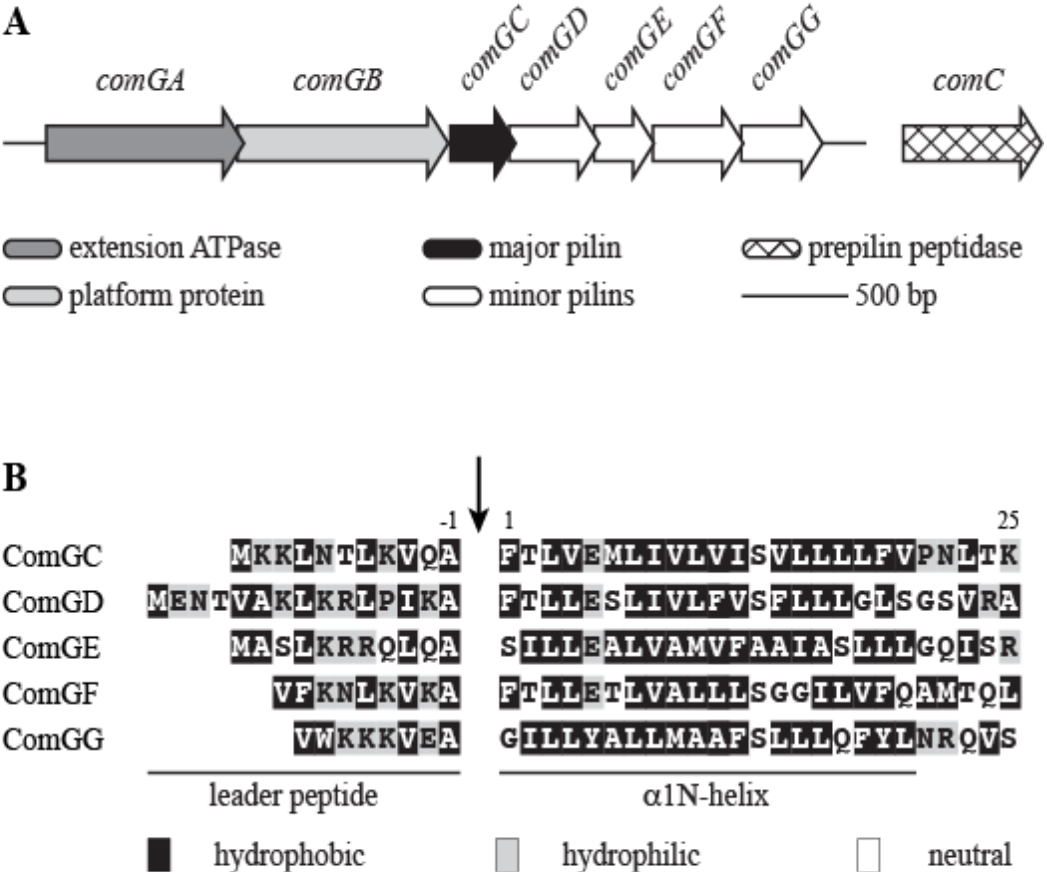
Com pilus machinery in *S. sanguinis*. (**A**) Genomic organisation of the genes involved in the biogenesis of the Com pilus in *S. sanguinis* 2908. All the genes are drawn to scale, with the scale bar representing 500 bp. The functions of the corresponding proteins are listed at the bottom. (**B**) Sequence alignment of the putative N-terminal class III signal peptides of the five ComG pilins in *S. sanguinis* 2908. The 8-15 aa-long leader peptides, which contain a majority of hydrophilic (shaded in grey) or neutral (no shading) residues, end with a conserved Ala-1. Leader peptides are processed (indicated by the vertical arrow) by the prepilin peptidase ComC. The mature proteins start with a tract of 21 predominantly hydrophobic residues (shaded in black), which invariably form the protruding N-terminal portion of an extended α-helix that is the main assembly interface within filaments.

We next determined the global distribution of the Com system in publicly available complete bacterial genomes using MacSyFinder (25). Specifically, we used the MacSyFinder model built for the identification of Com systems (2), which takes into account the genetic composition and organisation of its components. This showed that the Com system is restricted to Firmicutes, a phylum comprising a vast majority of monoderms, where it is exceptionally widespread since it was detected in 2,333 genomes (Supplemental Spreadsheet 1). An overwhelming majority of the corresponding species (99.7%) belong to the taxonomic class Bacilli (equally distributed among the Bacillales and Lactobacillales orders). As many as 88.7% of the sequenced Bacilli have a Com system. We also detected Com systems in one Clostridia (out of 336) and six Erysipelotrichia (out of 14). In total, 349 different species have the potential to express a Com pilus (Supplemental Spreadsheet 2).

Taken together, these findings suggest that the Com pilus is almost ubiquitous in Bacilli and can be advantageously studied in *S. sanguinis*.

### ComGC, the major subunit of Com pili, is highly conserved and defines a monophyletic group among type IV pilins

We next focused specifically on the major subunit of Com pili, the pilin ComGC (15,16). Compared to major pilins from T4aP, ComGC is ~40% shorter, with 94 or 93 aa for the processed ComGC_SS_ and ComGC_SP_, respectively (10.2 and 10.4 kDa). Moreover, unlike most other pilins, in which the only detectable sequence homology is usually in the α1N portion of the class III signal peptide (3), ComGC orthologs show extensive sequence identity. For example, processed ComGC_SS_ and ComGC_SP_ display 65.6% overall sequence identity (Fig. 2). Similarly, processed ComGC_SS_ and ComGC_BS_ (from *B. subtilis*) show 33.3% sequence identity overall (Fig. S1). This is consistent with the existence of a ComGC signature in the InterPro database (IPR016940) (4), which lists 2,809 ComGC entries. Global multiple alignment of these ComGC proteins shows that most of the sequence is conserved in ~90% of the entries (Fig. 2). In Fig. 2, the consensus sequences have been aligned to ComGC_SS_ and ComGC_SP_. Strikingly, some residues show sequence identity in virtually all the entries, including residues outside of the α1N portion (such as Ala_38_, Gln_46_, Tyr_50_ and Leu_64_ in ComGC_SS_).

**Fig. 2.**
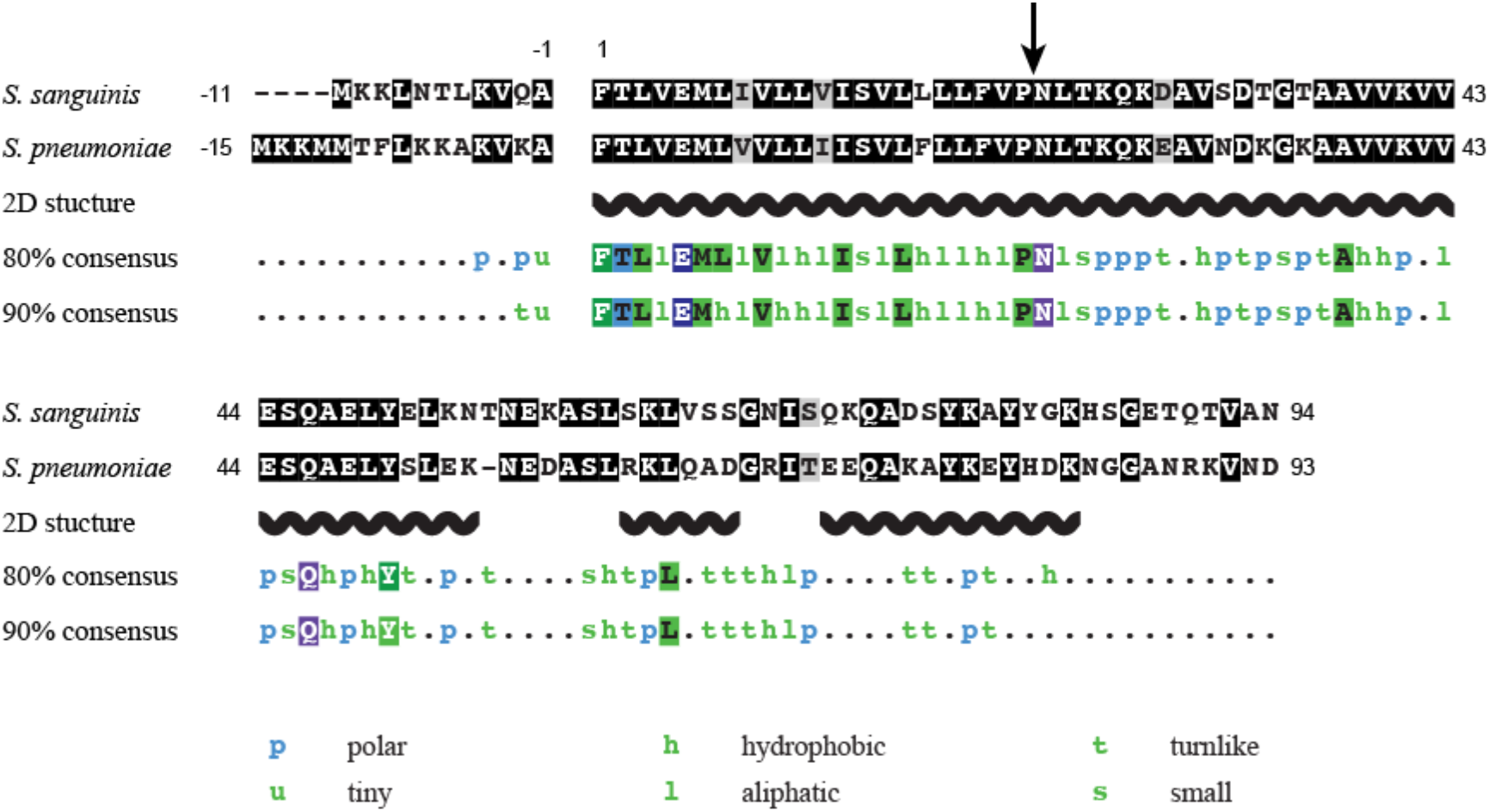
Global sequence analysis of ComGC pilins. Sequence alignments of ComGC in *S. sanguinis* and *S. pneumoniae* is represented in the top two rows. Residues were shaded in black (identical), grey (conserved) or unshaded (different). The leader peptide is highlighted. In the recombinant proteins that were produced for structure determination, the N-terminal 22 residues invariably forming a protruding hydrophobic α-helix were truncated (depicted by an arrow) to promote solubility. The 2D structural motifs predicted using JPred are depicted in the third row. Fourth and fifth rows represent the 80 and 90% ComGC consensus sequences, computed from 2,809 ComGC entries in InterPro, and aligned to ComGC_SS_ and ComGC_SP_. Multiple alignments were generated using Clustal Omega and formatted with MView. Polar: C, D, E, H, K, N, Q, R, S or T. Tiny: A or G. Hydrophobic: A, C, F, G, H, I, K, L, M, R, T, V, W or Y. Aliphatic: I, L or V. Turn-like: A, C, D, E, G, H, K, N, Q, R, S or T. Small: A, C, D, G, N, P, S, T or V.

The above observations suggest that Com pili form a highly homogeneous T4F subfamily. This was tested by performing a phylogenetic analysis based on the protein sequences of major pilins from different T4F found in a wide variety of bacteria, including T4aP, T4bP, T4cP (also known as Tad pili), mannose-sensitive hemagglutinin pili (MSH), type II secretion systems (T2SS) and Com pili. The phylogeny tree that was generated (Fig. 3), using IQ-TREE (26), reveals that several T4F are in clear monophyletic groups with good branch support, >96% ultrafast bootstrap (UFBoot) (27). Of particular interest, Com pili define a highly supported clade (99% UFBoot), clearly distinct form all other T4F systems.

**Fig. 3.**
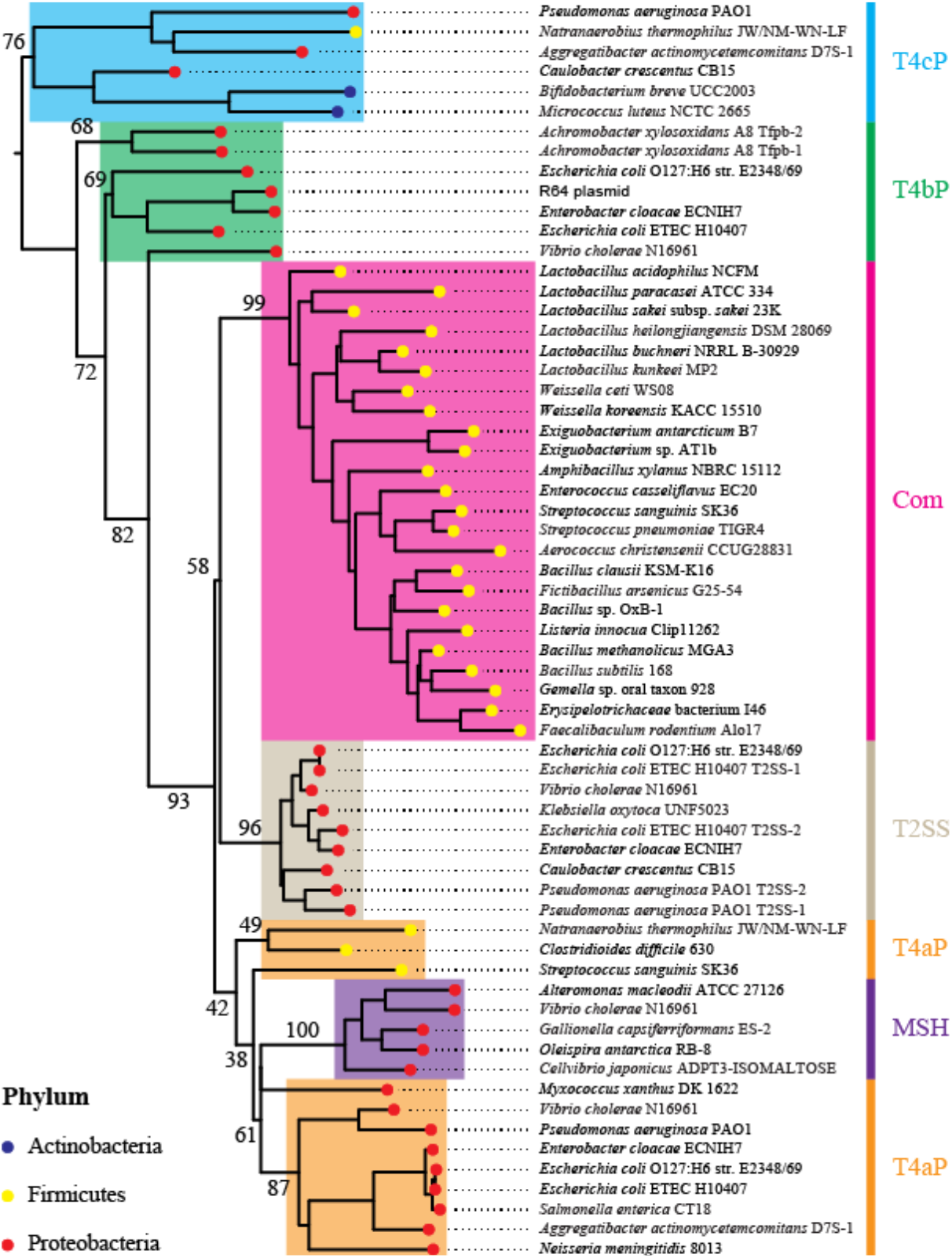
Rooted phylogeny of the major pilins from various bacterial T4F. The tree was build using IQ-Tree, with 1,000 replicates of UFBoot and LG+F+R4 model. Numeric values (in %) indicate UFBoot of the corresponding branches. The colour of the bullet points indicates the taxonomic group of the corresponding species. The colour of the strips and highlights indicate the classification of the different T4F systems. T4aP: type IVa pilus. T4bP: type IVb pilus. T4cP: type IVc pilus (also known as Tad). MSH: mannose-sensitive hemagglutinin pilus. Com: competence pilus. T2SS: type II secretion system.

Taken together, these findings show that ComGC is a small pilin with a highly conserved sequence, which defines a monophyletic group.

### Solution structure of two ComGC orthologs reveal a conserved and new type IV pilin fold

Since high-resolution structural information is needed to improve our understanding of Com pili, we decided to solve the 3D structure of ComGC_SS_. To facilitate protein purification, we used a synthetic *ComGC_SS_* gene codon-optimised for expression in *Escherichia coli*, and produced a soluble protein in which the first 22 residues of ComGC_SS_ that form a hydrophobic α-helix (α1N) were replaced by a non-cleavable N-terminal hexahistidine tag (6His). This commonly used truncation approach is predicted to have minimal structural impact on the rest of the protein, as previously shown for the *P. aeruginosa* PAK pilin (28). The resulting 8.8 kDa 6His-ComGC_SS_ protein could be readily purified using a combination of affinity and gel-filtration chromatography. After purification of isotopically labelled protein with ^13^C and 15N for backbone and side-chain NMR resonance assignments, we could assign 99.5% of the backbone and 92% of assignable protons overall. Structural ensembles were determined with 962 NOE based restraints, 50 hydrogen bonds, 110 dihedral angles restraints and 39 residual dipolar couplings (RDC) (Table 1). As can be seen in Fig. 4, ComGC_SS_ 3D structure is unlike that of any type IV pilin present in PDB, as it is purely helical, with three distinct helices connected by loops. The helices present are consistent with JPred secondary structure prediction (Fig. 2) (29). The N-terminal α1-helix, which involves residues 37-53 of the processed protein, corresponds to α1C since the hydrophobic α1N has been truncated in 6His-ComGC_SS_. Tightly packed against this α1-helix, in a parallel plane, are α2-helix (residues 61-67) and α3-helix (residues 72-85), which stack against each other in antiparallel fashion (Fig. 4A) and orthogonally to α1. Except for the N-terminal unstructured residues, the ComGC_SS_ structures within the NMR ensemble superpose well onto each other (Fig. 4B), with a root mean square deviation (RMSD) of 1.2 Å for Cα atoms, which suggests that there is no significant flexibility in this region of the structure (30). The unstructured N-terminus, which lacks long and medium NOEs present in the ordered regions of the proteins, was predicted to be highly dynamic based on TALOS+ (31), with an average S^2^ order parameter of 0.49 ± 0.10.

**Table 1.** NMR structural statistics.

|  | ComGC<sub>SS</sub> | ComGC<sub>SP</sub> |
| --- | --- | --- |
| **NOE-derived distance constraints** |  |  |
| long [(i-j) > 5] | 128 | 95 |
| medium [$5 \geq (i-j) > 1$] | 414 | 404 |
| intraresidue (i=j) | 420 | 381 |
| total | 962 | 880 |
| hydrogen bonds | 50 | 54 |
| dihedral constraints ($\Phi$ and $\Psi$) | 110 | 102 |
| residual dipolar couplings (RDC) | 39 | 38 |
| **Ramachandran statistics (from PROCHECK)** |  |  |
| most favoured (%) | 93.4 | 83.0 |
| additionally allowed (%) | 6.4 | 15.9 |
| generously allowed (%) | 0.2 | 1.1 |
| disallowed (%) | 0.0 | 0.0 |
| **Structure statistics** |  |  |
| RMSD backbone (all residues) | 3.3 | 4.0 |
| RMSD backbone (ordered residues*) | 0.6 | 0.8 |
| RMS bond angles ($^{\circ}$) | 1.8 | 1.9 |
| RMS bond lengths ($\text{\AA}$) | 0.012 | 0.017 |
| **Restraint statistics (RMSD of violations)** |  |  |
| NOE restraints | $0.060 \pm 0.003$ | $0.179 \pm 0.008$ |
| hydrogen bonds | $0.075 \pm 0.015$ | $0.100 \pm 0.017$ |
| dihedral restraints | $1.805 \pm 0.075$ | $1.827 \pm 0.318$ |
| RDC | $0.748 \pm 0.138$ | $0.716 \pm 0.256$ |
| Q value | $0.146 \pm 0.028$ | $0.150 \pm 0.054$ |

\*PROCHECK ordered residues are 37-53, 61-66 and 72-85 for ComGC<sub>SS</sub>, and 36-54, 60-65 and 71-82 for ComGC<sub>SP</sub>.

**Fig. 4.**
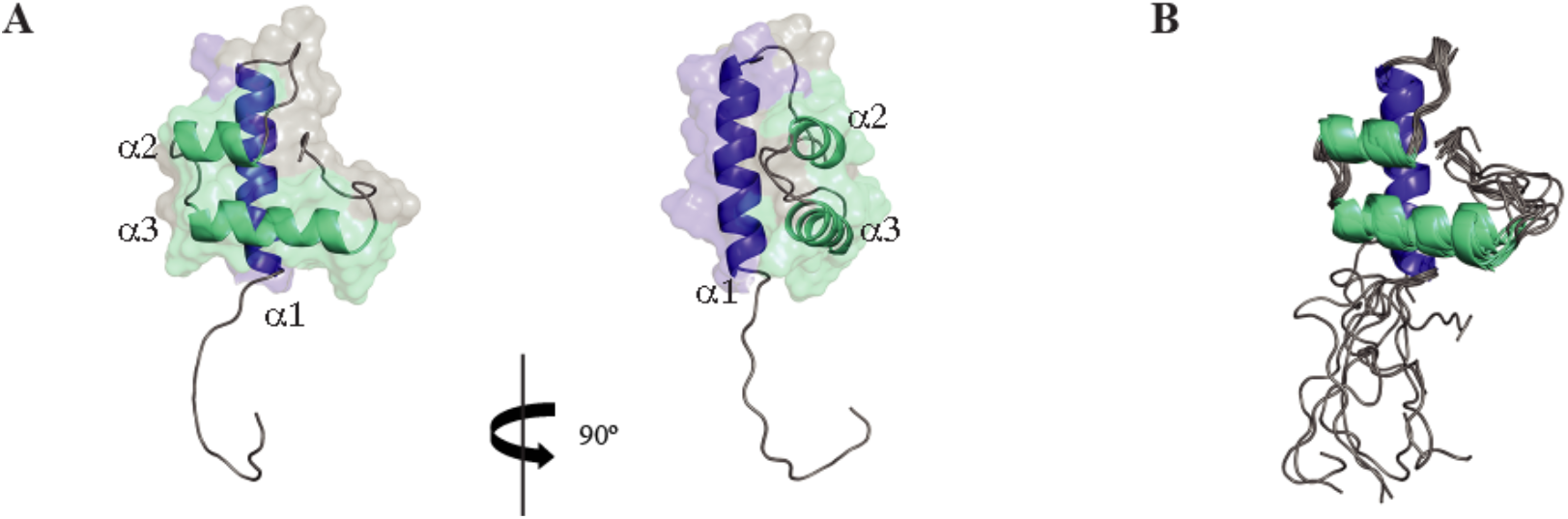
3D solution structure of ComGC_SS_. (**A**) Cartoon representation of the ComGC_SS_ structure: face and side views are shown. A dimmed surface representation of the protein is superimposed. The three consecutive α-helices have been named α1, α2 and α3, and highlighted in blue (α1) or cyan (α2 and α3). (**B**) Cartoon representation of the superposition of the ensemble of 10 ComGC_SS_ structures determined by NMR, which highlights that there is no significant flexibility in the structure except for the unstructured N-terminus.

Our ComGC_SS_ structure differs markedly from the recently reported solution structure of ComGC_SP_ (PDB 5NCA) (19), which is surprising considering the high sequence identity between these two proteins (Fig. 2). Therefore, in order to define the structural relationship between ComGC orthologs, we decided to solve the structure of ComGC_SP_. As above, we used a synthetic *ComGC_SP_* gene codon-optimised for expression in *E. coli*, we fused the 71 aa-long soluble portion of ComGC_SP_ to a non-cleavable N-terminal 6His tag and purified doubly labelled 6His-ComGC_SP_ (9 kDa). Again, assignment was excellent since 98.1% of the backbone and 90% of assignable protons overall could be assigned. Structural ensembles were determined with 880 NOE based restraints, 54 hydrogen bonds, 102 dihedral angles restraints and 38 RDC (Table 1). As can be seen in Fig. 5, our ComGC_SP_ 3D structure is highly similar to the structure of ComGC_SS_. It is, however, very different (RMSD of 3.6 Å) from the ComGC_SP_ solution structure that was recently determined from a low number of restraints (Fig. S2) (19). In brief, our structure shows that ComGC_SP_ displays three distinct helices, with α2-helix (residues 60-66) and α3-helix (residues 71-82) stacking against each other and packing orthogonal to the N-terminal α1-helix (Fig. 5A). As for ComGC_SS_, except for the unstructured N-terminus, there is no significant flexibility in ComGC_SP_ since the structures within the NMR ensemble superpose well onto each other, with a RMSD of 1.6 Å for Cα atoms (Fig. 5B). Our ComGC_SS_ and ComGC_SP_ averaged structures are highly similar (Fig. 5C), with 1.8 Å RMSD between their ordered regions and 1.5 Å RMSD for the helical regions, which is consistent with the high sequence identity between these two proteins.

**Fig. 5.**
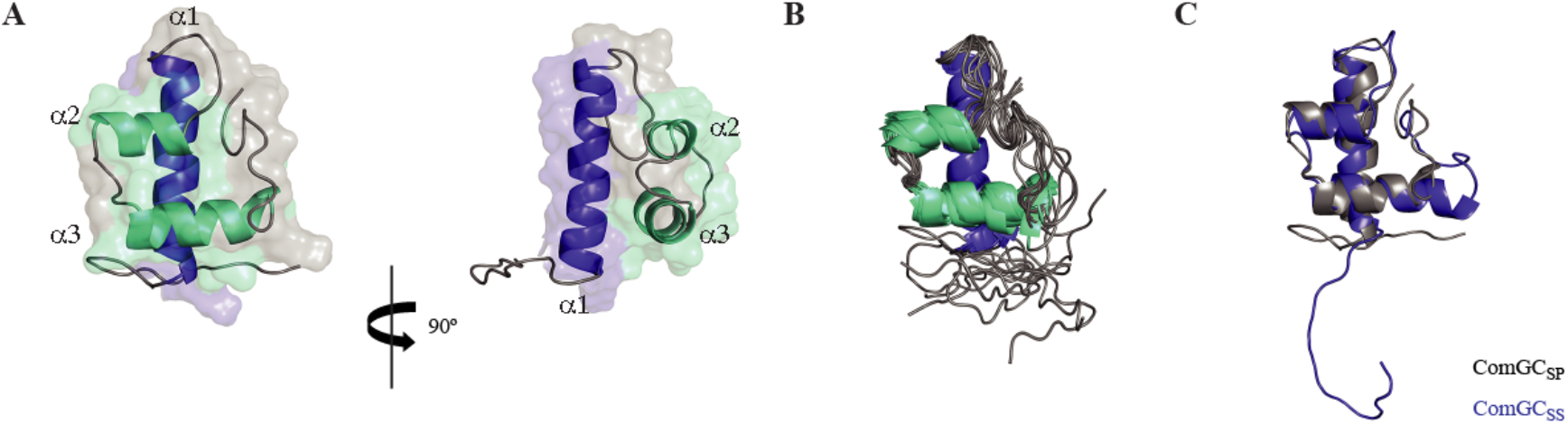
3D solution structure of ComGC_SP_. (**A**) Cartoon representation of the ComGC_SP_ structure: face and side views are shown. A dimmed surface representation of the protein is superimposed. Nomenclature and colour scheme are the same than in Fig. 4. (**B**) Cartoon representation of the superposition of the ensemble of 10 ComGC_SP_ structures determined by NMR, (**C**) Cartoon representation of the overlay of ComGC_SP_ and ComGC_SS_ representative structures. This highlights the high structural similarity between the two proteins, with 1.5 Å RMSD for the helical regions.

As determined by GETAREA (32) with a probe radius of 1.4 Å, the average ratio of solvent exposure for the ordered portion of ComGC_SS_ is 48.3%, relative to 6.7% for those residues determined to be on the interior. In our ComGC_SS_ structure, conserved residues Val_43_, Gln_46_, Tyr_50_, Leu_64_ and Ile_70_ are deeply buried, with an average of only 6% solvent exposure, forming a critical portion of a hydrophobic core contributing to the globular fold of ComGC. (Fig. 6). In contrast, conserved Gly_68_ is solvent exposed, which is important for the formation of the α2-helix-turn-α3-helix motif where a tiny residue at the beginning of the turn is necessary to provide the flexibility and lack of steric restrictions required for turning. These observations also apply to our ComGC_SP_ structure and are surprisingly reflected in the conservation of multiple chemical shifts between the conserved residues in our two structures (Fig. S3). In addition, modelling of the globular head of ComGC_BS_ (Fig. S4), which predicts a globular fold similar to ComGC_SS_ and ComGC_SP_, shows that Cys36 and Cys76 are in close enough proximity to form a disulfide bond. Such disulfide bond, which is absent in ComGC_SS_ and ComGC_SP_ that do not have Cys residues, is expected to stabilise the globular fold and was reported to stabilise ComGC in *B. subtilis* (16,33).

**Fig. 6.**
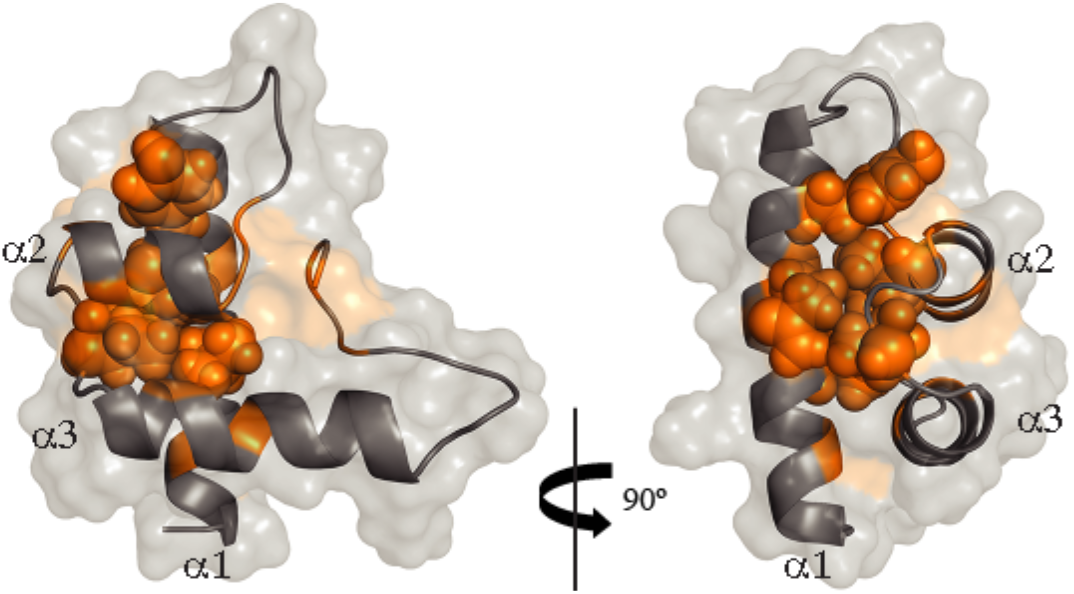
Conserved residues contributing to the globular fold of ComGC. Cartoon representation of the ordered portion of ComGC_SS_, where residues determined to be on the interior using GETAREA – with accessible surface accessibility ratios of less than 20% -are highlighted in orange. The consensus residues Val_43_, Gln_46_, Tyr_50_, Leu_64_ and Ile_70_ are shown with space filling representation.

Since the hydrophobic α1N that has been truncated in 6His-ComGC_SS_ is highly similar to the corresponding portion of several other bacterial T4F major pilins, including the PilE major pilin from *Neisseria gonorrhoeae* (Fig. S5) for which a full-length crystal structure is available (34), we could model the structure of the portion of α1 truncated in our construct to produce a full-length model of ComGC_SS_ (Fig. 7). Comparison with the two different pilin folds identified so far – pilins from *N. gonorrhoeae* and *Geobacter sulfurreducens* have been chosen as representative models – clearly shows that ComGC adopts a radically different type IV pilin fold (Fig. 7). All three pilins have in common an extended N-terminal α1-helix, the universal defining structural feature of type IV pilins (3). In addition, while the very short *G. sulfurreducens* pilin almost exclusively consists of α1, both ComGC and PilE display a typical lollipop shape with a globular head mounted onto a “stick” (the α1-helix). However, unlike in canonical pilins where the globular head consists of a 4-to 7-stranded antiparallel β-sheet in a parallel plane to α1, oriented 45° or more relative to the long axis of α1 (3), in ComGC the structural backbone of the globular head is an helix-turn-helix roughly orthogonal to α1 (Fig. 7). This fold, which falls within the class of mainly α and the architecture of orthogonal bundles, represents a novel pilin fold.

**Fig. 7.**
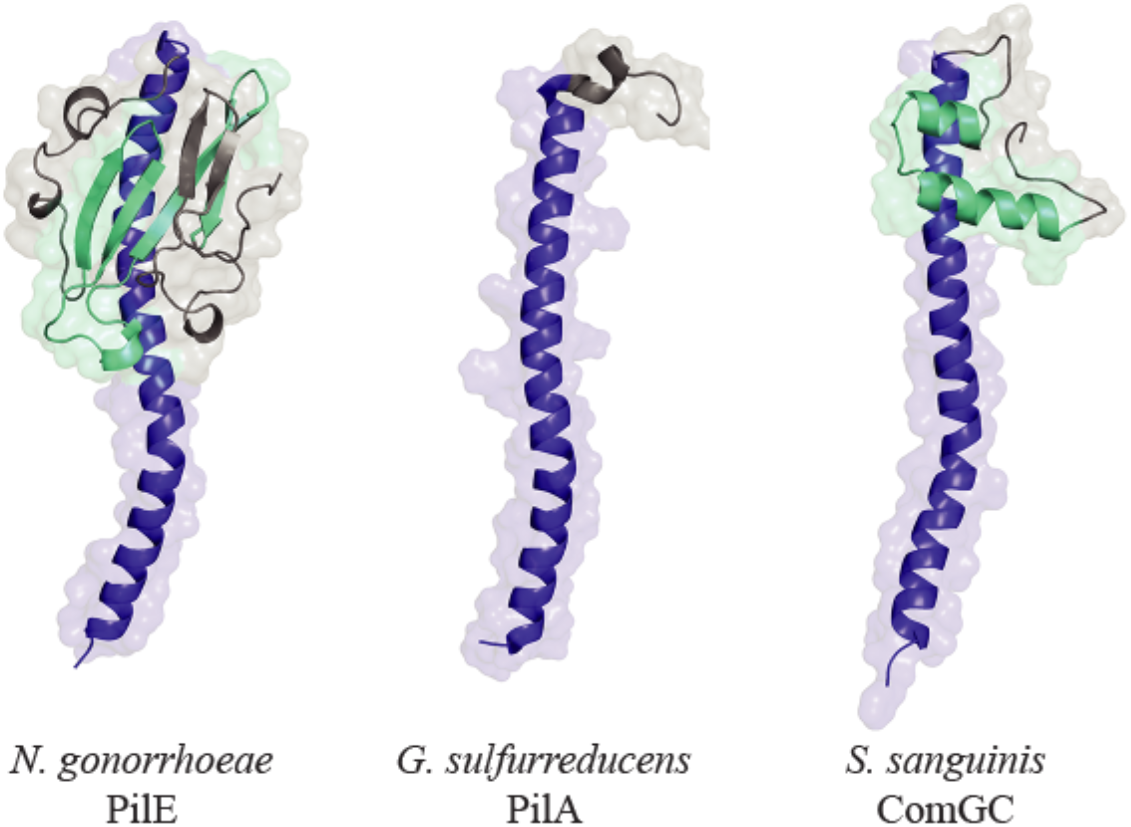
ComGC display a novel type IV pilin fold. 3D structure of the three different structural types of type IV pilins identified so far. The canonical type IV pilin fold is represented by the major pilin of T4aP in *N. gonorrhoeae* (PDB 2PIL). *G. sulfurreducens* T4P pilin (PDB 2M7G) is the chosen representative of the very short pilins almost exclusively consisting of α1. The full-length 3D structure of ComGC_SS_ has been modelled. The conserved α1 is highlighted in blue. Distinctive structural features in the globular heads of PilE (antiparallel β-sheet) and ComGC (antiparallel α2-α3 orthogonal to α1) have been highlighted in cyan.

Taken together, these structural findings show that ComGC orthologs display conserved 3D structures, with a previously unreported type IV pilin fold.

### ComGC novel pilin fold is compatible with helical T4F assembly

Since ComGC represents a novel type IV pilin structural fold, it was important to determine whether it could be modelled into recent cryo-EM structures obtained for a variety of bacterial T4F (7,8,35). These similar structures, *i.e.* filaments are right-handed helical polymers where pilins are held together by interactions between their α1 helices within the filament core, have revealed that a segment of α1N is melted during filament assembly, centred on helix-breaking residues Pro22. That portion of α1 is highly conserved in ComGC, including the helix-breaking Pro22 (Fig. S5). Using SWISS-MODEL (36) and the cryo-EM structure of *N. gonorrhoeae* T4P (8) as a template, we produced a full-length 3D structural model of ComGC_SS_ with a melted α1N segment (Fig. 8A). Considering that ComGC defines a monophyletic group and is highly conserved, it is very likely that all ComGC orthologs will display a similar 3D structure. This notion was strengthened by producing structural models for a range of different species expressing more or less distant ComGC (21.3-65.6% sequence identity), which were used to generate the phylogeny tree in Fig. 3. As seen in Fig. S6, all the models display the same lollipop shape with a globular head mounted onto a α1 stick. As for ComGC_SS_ and ComGC_SP_, the structural backbone of the globular head is always a helix-turn-helix roughly orthogonal to α1.

**Fig. 8.**
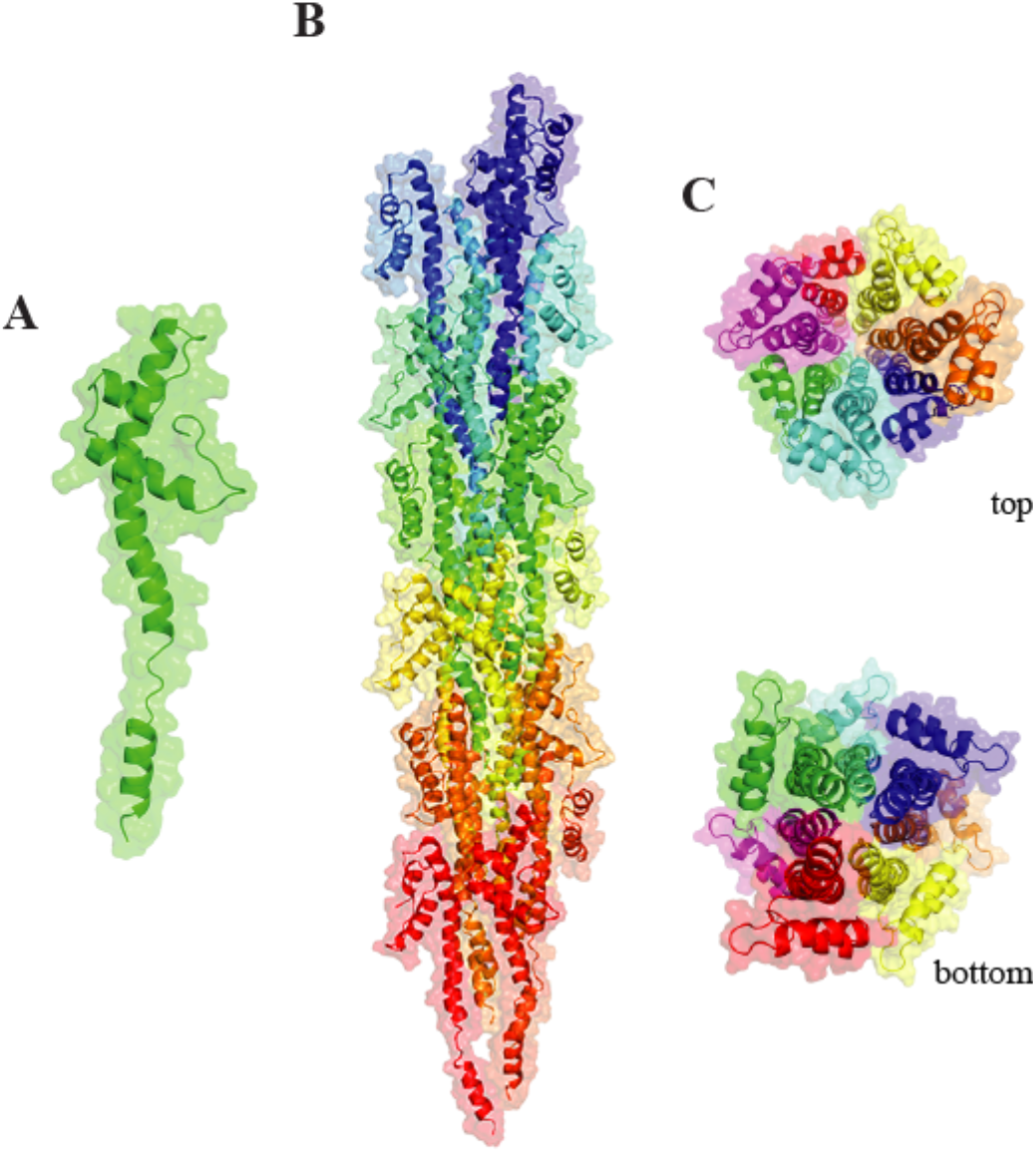
3D model of ComGC filaments. The cryo-EM structure of the *N. gonorrhoeae* T4P (PDB 5VXX) has been used as a template to generate a model of ComGC_SS_ pili. (**A**) Full-length ComGC_SS_ in filaments with a melted segment in α1N. (**B**) ComGC_SS_ pili with a right-handed helical packing of the conserved α1-helices which run approximately parallel to each other in the filament core. (**C**) Top and bottom views of ComGC_SS_ pili highlighting the globular heads forming the outer surface of the filaments and the extensive interactions between α1-helices in the filament core.

We next assessed whether full-length ComGC would be compatible with helical T4F assembly and found that to be the case. Despite its novel pilin fold, we were able to model packing of ComGC within the cryo-EM structure of *N. gonorrhoeae* T4P (8). This produced a homology model with good Ramachandran plot statistics based on PROCHECK (37), *i.e.* allowed (89.5%), additional allowed (8.2%), generously allowed (2.3%) and disallowed (0%). As can be seen in Fig. 8B, the model revealed a right-handed helical packing of the conserved N-terminal α1-helices of ComGC_SS_ in the filament core, which run approximately parallel to each other and establish extensive hydrophobic interactions (Fig. 8C). In addition, the Glu5 side chain of subunit S establishes a salt bridge and a hydrogen bond with Phe_1_ and Thr_2_, respectively, of S+1. Importantly, the globular heads are stacked on top of each other along the long axis of the filaments and their helix-turn-helix structural backbone forms the outer surface of the filaments (Fig. 8C). A very similar model was obtained using *P. aeruginosa* T4P as a template (8).

## Discussion

Their virtual ubiquity in prokaryotes and role in a variety of key biological processes make T4F an important research topic (1,2). Com pili are involved in DNA uptake in naturally competent monoderm bacteria (9). Imported DNA, which usually leads to genome diversification via transformation, can also be used as a source of food or as a template for repair of damaged genomic DNA (10). Compared to T4F in diderms, most notably T4P and T2SS that have been extensively studied, Com pili have been understudied, including from a structural point of view. In this report, we focused on the major subunit of Com pili, the ComGC pilin, which we analysed genomically, phylogenetically and structurally. This led to the notable findings discussed below.

Although Com pili have been primarily studied in two model competent species (*B. subtilis* and *S. pneumoniae*), the present study suggests that they are widespread since complete sets of Com pilus-encoding genes are readily detected in more than 2,300 genomes corresponding to almost 350 different species. However, unlike promiscuous T4F such as T4aP and T4cP that are found in virtually all phyla of Bacteria (2), Com pili are restricted to a single phylum (Firmicutes) and almost exclusively to a single underlying class of monoderms (Bacilli), where they are virtually ubiquitous. Indeed, an overwhelming majority of Bacilli genomes (88%) have Com-encoding genes. Interestingly, the major subunit of Com pili (ComGC) shows extensive sequence conservation in the corresponding genomes and define a clear monophyletic group within type IV pilins. Taken together, these observations suggest that the Com pilus is a T4F that has emerged only once, very early during the diversification of Firmicutes, where it has remained largely confined ever since. Since the Com-encoding genes have not become pseudogenes, it is likely that most Bacilli have the ability to assemble a Com pilus and take up DNA. However, since only a handful of these species have been experimentally shown to be competent (10), this implies that either the imported DNA is primarily used as food or for genome repair instead of genome diversification, or that the inducing cues leading to transformation are yet to be established for most species of Firmicutes. Alternatively, Com pili might have evolved in some of these species to take up other macromolecules, which is however at odds with the conservation of the five pilins.

Perhaps the most important finding in this study is that ComGC, the major subunit of the Com pilus, displays an entirely novel major pilin fold where the extended N-terminal α1-helix, the universal defining structural feature of type IV pilins (3), is topped by a purely helical globular head. ComGC thus appears to be a “middle ground” between longer canonical pilins (e.g. *N. gonorrhoeae*), in which the globular head consists of an antiparallel β-sheet, and the very short pilins where a globular head is missing altogether (e.g. *G. sulfurreducens*). These structures point to a hypothetical evolutionary scenario during which truncation of the antiparallel β-sheet in a canonical type IV pilin might have led to a purely helical ComGC proto-structure. Intriguingly, this scenario “works” particularly well with PilE1, a major subunit of *S. sanguinis* T4P, which has two short α-helices in the loop connecting α1 and the antiparallel β-sheet (11). Importantly, this putative “truncation” would not interfere with the expected ability of ComGC to be assembled into helical filaments, since this pilin could be readily modelled into recent T4F structures (7,8,35). Com pili are thus likely to result from the right-handed helical packing of ComGC α1-helices within the filament core, running parallel to each other and establishing extensive hydrophobic interactions, with a melted central portion. Such packing will stack the globular heads on top of each other, forming the surface of the filaments. Extensive sequence conservation, including for residues beyond the classically conserved α1N, and the fact that the two structures that we have solved are virtually identical, strongly suggest that these structural features apply to the whole ComGC clade, including species such as *B. subtilis* where extended filaments have not been observed (16). It is therefore surprising that a recently published NMR structure of ComGC_SP_ (PDB 5NCA) (19) differs dramatically from ours. While the previous structure is purely helical as well, the orientation of the α2 and α3 helices is entirely different, resulting in an absence of packing of the conserved hydrophobic core. Therefore, PDB 5NCA which resembles a one-sided “pick-axe” with no globular head cannot be readily modelled into recent T4F structures (Fig. S7). Interestingly, our assignments vary only slightly from those previously produced for PDB 5NCA (Fig. S8). However, while we have managed to successfully assign 90% assignable protons overall, the previous assignment was merely 65% (19), which probably accounts for the apparently “unfolded” state of PDB 5NCA. Indeed, without a high degree of proton identification, the assignment of NOESY peaks and production of distance restraints fails. Local hydrogen bonds and dihedral restraints often cannot compensate for lack of long-range NOEs within the protein interior or between elements of secondary structure.

Together with these conserved structural features, the conservation *en bloc* of the genes encoding the Com pilus strongly suggests that the molecular mechanisms of filament assembly and DNA uptake are widely conserved in Firmicutes. These mechanisms, which remain poorly understood, can be advantageously studied in *S. sanguinis*, which has recently emerged as a monoderm T4P model (20). Actually, *S. sanguinis* is so far the only monoderm likely to express two distinct T4F, Com pili and retractable T4aP, which further cements it as a prime model species. Comparison with other T4F systems shows that the machinery involved in biogenesis of Com pili is one of the simplest, by far. Since ComGD, ComGE, ComGF and ComGG pilins are likely to be minor pilus components important for filament stability and function (a conserved role for minor pilins in various T4F) (1), and ComC is the prepilin peptidase processing pilins (38), it appears that assembly of ComGC into filaments is mediated by two proteins only. Namely, an extension ATPase (ComGA) and a platform protein (ComGB), which together will assemble processed ComGC into a right-handed helical filament. Upon DNA binding, which has been visualised for *S. pneumoniae* Com pili, but the receptor is yet to be identified (15), uptake will be initiated by filament retraction (14). Since there is no dedicated retraction ATPase, one possibility is that ComGA might be a bifunctional motor powering both extension and retraction like recently suggested for the T4cP motor (39). It would be interesting to image Com filaments dynamics and DNA-binding ability in live cells, using a labelling strategy that has recently enabled the visualisation of these steps for T4aP involved in competence in naturally competent diderm species (14).

In conclusion, by providing high-resolution structural information for the ComGC pilins, this study has shed light on an understudied T4F involved in DNA uptake found in hundreds of monoderm bacterial species and has led to the surprising discovery of a novel type IV pilin fold. This paves the way for further investigations of this minimalist T4F, which are expected to improve our understanding of a fascinating superfamily of filamentous nanomachines ubiquitous in prokaryotes.

## Experimental procedures

### Bioinformatic analyses

Protein sequences were routinely analysed using the DNA Strider program. Protein sequence alignments were done using the Clustal Omega server at EMBL-EBI. Pretty-printing of alignment files was done using BoxShade server at ExPASy. Reformatting of large multiple alignment files was done using the MView server at EMBL-EBI. Prediction of functional domains was done using the InterProScan server at EMBL-EBI, which was also used to download all the ComGC protein entries with an IPR0160940 domain. Protein secondary structure prediction was done using JPred server at University of Dundee. Protein 3D structures were downloaded from the RCSB PDB server. Molecular visualisation of protein 3D structures was done using PyMOL (Schrödinger). The GETAREA server, at UTMB, was used for calculating the solvent accessible surface area of ComGC proteins.

Detection of the Com systems in genomes available in NCBI RefSeq database (last accessed in April 2019, 13,512 genomes of Bacteria and Archaea) was done as described previously (2), using MacSyFinder (25) and the relevant HMM Com model (2). Phylogenetic analysis based on protein sequences of major pilins of different T4F involved an initial alignment of the sequences using MAFFT v7.273 (40), specifically the linsi algorithm. Multiple alignments were analysed using Noisy v1.5.12 (41) with default parameters, in order to select the informative sites. Next, we inferred maximum likelihood trees from the curated alignments using IQ-TREE v 1.6.7.2 (26), with option -allnni. We evaluated the node supports using the options -bb 1,000 for ultra-fast bootstraps, and -alrt 1,000 for SH-aLRT (27). The best evolutionary model was selected with ModelFinder (42), option -MF and BIC criterion. We used the option -wbtl to conserve all optimal trees and their branches length.

### Protein expression and purification

A synthetic gene, codon-optimised for *E. coli* expression, encoding ComGC_SS_ from *S. sanguinis* 2908 (21) was synthesised and cloned by GeneArt, yielding pMA-T-*ComGC_SS_* (Table S2). The portion of the gene encoding residues 23-94 from the mature protein was PCR-amplified using *ComGC_SS_*-F and *ComGC_SS_*-R primers (Table S3), cut with NcoI and BamHI and cloned into the pET28b vector (Novagen) cut with the same enzymes. The forward primer was designed to fuse a non-cleavable N-terminal 6His tag to ComGC_SS_. The resulting plasmid was verified by sequencing and transformed into chemically competent *E. coli* BL21(DE3) cells. A single colony was transferred to 10 ml of LB supplemented with 50 μg.ml^−1^ kanamycin and grown at 37°C overnight (O/N). This pre-culture was back-diluted 100-fold into 1 l M9 minimal medium, supplemented with antibiotic, a mixture of vitamins and trace elements, and ^13^C D-glucose and ^15^N NH4Cl for isotopic labelling. Cells were grown in an orbital shaker at 37°C until the OD_600_ reached 0.7, before adding 0.4 mM IPTG (Merck Chemicals) to induce protein expression during 16 h at 18°C. Cells were then harvested by centrifugation at 8,000 *g* for 20 min and subjected to one freeze/thaw cycle in lysis buffer (PBS pH 7.4, with EDTA-free protease inhibitors). This lysate was further disrupted by repeated cycles of sonication, pulses of 5 sec on and 5 sec off during 5 min, until the cell suspension was visibly less viscous. The cell lysate was then centrifuged for 20 min at 18,000 *g* to remove cell debris. The clarified lysate was then passed using an ÄKTA Purifier FPLC through a 1 ml HisTrap HP column (GE Healthcare), pre-equilibrated in lysis buffer. The column was then washed extensively with lysis buffer to remove unbound material before 6His-ComGC_SS_ was eluted using elution buffer (PBS pH 7.4, 200 mM NaCl, 300 mM imidazole). Affinity-purified ComGC_SS_ was further purified by gel-filtration chromatography on an HiLoad 16/600 Superdex 75 column (GE Healthcare), using (25 mM Na_2_HPO_4_/NaH_2_PO_4_ pH 6, 200 mM NaCl) buffer for elution. For RDC measurements we produced ^15^N labelled protein as follows. Bacteria grown O/N in 5 ml LB with antibiotic were sub-cultured at 37°C in 0.8 l LB to 0.6 OD_600_, and then transferred to 0.4 l M9 with ^15^N NH_4_Cl, unlabelled D-glucose, and 10 μg.l^−1^ thiamine. Cultures were induced with 0.3 mM IPTG at 16°C for 18 h. After the production of a clarified lysate, protein was purified as above, except for the use of hand-made Ni-NTA agarose (Qiagen) in (50 mM Tris pH 8, 300 mM NaCl) and eluted using (50 mM Tris pH 8, 200 mM NaCl, 300 mM imidazole), and Superdex 75 10/300 GL (GE Healthcare) columns in (25 mM Tris pH 8, 200 mM NaCl) and dialysed into (25 mM Na_2_HPO_4_/NaH_2_PO_4_ pH 6, 50 mM NaCl).

For ComGC_SP_, a codon-optimised synthetic gene based on the gene from *S. pneumoniae* R6 was synthesised and cloned by GeneArt, yielding pMA-T-*ComGC_SP_* (Table S2). The portion of the gene encoding residues 23-93 from the mature protein was PCR-amplified using *ComGC_SP_*-F and *ComGC_SP_*-R primers (Table S3), cut with NcoI and BamHI and cloned into the pET28b vector (Novagen) cut with the same enzymes. The forward primer was designed to fuse a non-cleavable N-terminal 6His tag to ComGC_SS_. The resulting plasmid was verified by sequencing and transformed into chemically competent *E. coli* BL21(DE3) cells. A single colony was transferred to 5 ml of LB supplemented with 50 μg.ml^−1^ kanamycin and grown O/N at 37°C. Bacteria were sub-cultured at 37°C in 0.8 l LB with antibiotic to OD_600_ 0.7, and then transferred into 0.4 l M9 with10 μg.l^−1^ thiamine, and either ^15^N NH4Cl and unlabelled D-glucose, or ^15^N NH4Cl and ^13^C D-glucose. Cultures were induced with 0.3 mM IPTG at 16°C for 18 h. After the production of a clarified lysate, ComGC_SS_ was purified as above using hand-made Ni-NTA agarose (Qiagen) and Superdex 75 10/300 GL (GE Healthcare) columns.

### NMR spectroscopy and structure determination

All data was collected on Bruker Avance III HD 800 MHz and 600 MHz triple resonance spectrometers with cryoprobes operated at 25°C. For ComGC_SS_, a sample containing ^13^C, ^15^N labelled protein at 1 mM in NMR buffer (25 mM Na_2_HPO_4_/NaH_2_PO_4_ pH 6, 50 mM NaCl, 5% D_2_O) was used for assignment experiments and structure determination. For ComGC_SP_, a sample containing ^13^C, 15N labelled protein at 1.8 mM in NMR buffer was used for assignment experiments and structure determination. Resonance assignments for ComGC_SS_ were performed using ^15^N HSQC, ^13^C aliphatic HSQC, HNCACB, CBCACONH, HBHA, HNCO, HNCACO, HCCCONH, CCCONH and CCH. For ComGC_SP_, assignments were performed using ^15^N HSQC, ^13^C aliphatic HSQC, HNCA, CBCANH, CBCACONH, HBHA, HNCO, HNCACO, HCCCONH, CCCONH and CCH. All data was processed using MddNMR (43) for reconstruction after Non-Uniform Sampling and NMRPipe (44). Peak picking and assignments were performed in SPARKY (45).

NOE peak lists were used, with mixing time of 140 msec, from 3D ^13^C HSQC-NOESY, 3D ^15^N HSQC-NOESY for ComGC_SP_, and simultaneous ^13^C/^15^N chemical shift evolution NOESY for ComGC_SS_. For both proteins, RDC lists were derived from ^15^N HSQC-IPAP experiments on ^15^N labelled isotropic and aligned sample in 3% PEG/hexanol liquid crystal, with D_2_O splitting of ~7 Hz. RDCs were included in the structure calculations if there was baseline resolution and for residues where TALOS+ predicted order parameter of >0.8. Angular constraints from TALOS+ were used in the structure calculations. Both ComGC_SS_ and ComGC_SP_ structures were determined using Ponderosa-C/S (45), refined using Xplor-NIH 2.52 (46), aligned using Theseus (47), and secondary structure checked using Stride (48). Structure validation was performed using PSVS (49), PROCHECK (37) and in-house scripts.

### Modelling

SWISS-MODEL server at ExPASy was used for modelling protein 3D structures. In brief, the full-length ComGC_SS_ was modelled with using *N. gonorrhoeae* major pilin PilE (PDB 2PIL) as a template (50). We first modelled the missing α1 residues in our structure, which was aligned to our Xplor-NIH-produced average NMR structure (without the first unstructured α1 residues) using PyMOL and finally merged using Coot (51).

Similarly, the full-length ComGC_SS_ structure within filaments was modelled by using one of the PilE subunits from the cryo-EM model of <u>*N. gonorrhoeae*</u> T4P (PDB 5VXX) (8) as a template for the missing α1 residues in our structure. The Com pilus model was produced after alignment of the averaged NMR structure ComGC_SS_ α1-helices to the α1-helices of SWISS-MODEL PilE-based homology model subunits in the <u>*N. gonorrhoeae*</u> T4P. This was also done for the recently published ComGC_SP_ structure (PDB 5NCA). The structural elements were fused using Coot (51). In addition, we modelled packing of full-length ComGC_SS_ in the PAK pilus from *P. aeruginosa* (PDB 5VXY) (8).

## Acknowledgments

This work relied heavily on the use of the Cross-Faculty NMR Centre at Imperial College London. We are grateful to Nicolas Biais (City University of New York) and Romé Voulhoux (CNRS Marseille) for critical reading of the manuscript.

## Data availability

The NMR solution structures of ComGC_SS_ and ComGC_SP_ have been deposited in the Protein Data Bank under accession numbers 6TXT and 6Y1H, respectively. Chemical shift assignments and NOE-based restraints used in structure calculations are available from the Biological Magnetic Resonance Data Bank under accession numbers 34477 and 34490, respectively. All the other data described in the manuscript are either contained within the manuscript, or are to be shared upon request to corresponding author Vladimir Pelicic.

## Footnotes

This work was funded by a Medical Research Council grant (MR/P022197/1) to V.P. R.D. was funded by the doctoral school Complexité du vivant-ED515 (contract number 2449/2016). E.P.C.R. was funded by the INCEPTION project (PIA/ANR-16-CONV-0005).

## The abbreviations used are

T4F: type IV filaments
T4P: type IV pili
Com: competence
UFBoot: ultrafast bootstrap
NMR: nuclear magnetic resonance
NOE: nuclear Overhauser effect
RDC: residual dipolar couplings
RMSD: root mean square deviation
PDB: Protein Data Bank
LB: Lysogenic Broth
O/N: overnight
PBS: phosphate-buffered saline
IPTG: isopropyl β-D-1-thiogalactopyranoside
EDTA: ethylenediaminetetraacetic acid

