## Supplemental information for "The major subunit of widespread competence pili exhibits a novel and conserved type IV pilin fold"

**Table S1. Bioinformatic analysis of the components of *S. sanguinis* Com machinery.**

| Name | MW (Da)* | pI* | InterPro domain | predicted function | identity in *B. subtilis* (%)* | identity in *S. pneumoniae* (%)* |
|---|---|---|---|---|---|---|
| ComGA | 35,864 | 6.17 | IPR001482 | extension ATPase | 33.01 | 74.76 |
| ComGB | 37,817 | 9.06 | IPR003004 | platform protein | 21.36 | 67.37 |
| ComGC | 10,184 | 7.70 | IPR016940 | major pilin | 33.33 | 65.59 |
| ComGD | 14,680 | 6.40 | - | minor pilin | 20.66 | 55.04 |
| ComGE | 9,689 | 10.17 | IPR021749 | minor pilin | 13.92 | 52.33 |
| ComGF | 15,787 | 5.25 | IPR016977 | minor pilin | 24.41 | 61.76 |
| ComGG | 13,663 | 9.32 | - | minor pilin | 19.80 | 38.84 |
| ComC | 24,816 | 9.23 | IPR010627 | prepilin peptidase | 24.20 | 57.53 |

*For the pilins, the numbers correspond to the processed proteins.

**Table S2. Plasmids used in this study.**

| **Name** | **Description** | **Source** |
|---|---|---|
| pET28-b | T7-based expression vector | Novagen |
| pMA-T-*comGC*$_{SS}$ | pMA-T-derivative with codon-optimised *comGC*$_{SS}$ | this study |
| pMA-T-*comGC*$_{SP}$ | pMA-T-derivative with codon-optimised *comGC*$_{SP}$ | this study |
| pET28-*comGC*$_{SS}$ | pET28-derivative for expressing $His_6$-ComGC$_{SS}$ | this study |
| pET28-comGC$_{SS}$ | pET28-derivative for expressing $His_6$-ComGC$_{SP}$ | this study |

**Table S3. Primers used in this study.**

| Name | Sequence* |
|---|---|
| *comGC*$_{SS}$-F | ggg**ccatgg**atcatcatcatcatcatcatAATCTGACCAAACAGAAAGATGCAGTTAGC |
| *comGC*$_{SS}$-R | ccc**ggatcc**TTAATTTGCAACGGTCTGGGTTTCACCG |
| *comGC*$_{SP}$-F | ggg**ccatgg**atcatcatcatcatcatcatAATCTGACCAAACAGAAAGAAG |
| *comGC*$_{SP}$-R | ccc**ggatcc**TTAATCGTTCACTTTGCGATTGGCACC |

*Overhangs are in lower case. Restriction sites are in bold.

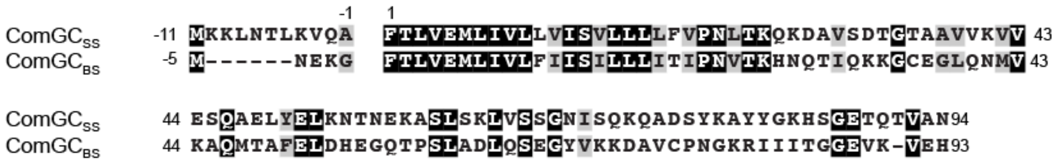


**Fig. S1. Sequence alignment of ComGC in *S. sanguinis* and *B. subtilis.*** Residues were shaded in black (identical), grey (conserved) or unshaded (different). The leader peptide, which is processed by the prepilin peptidase ComC is highlighted.

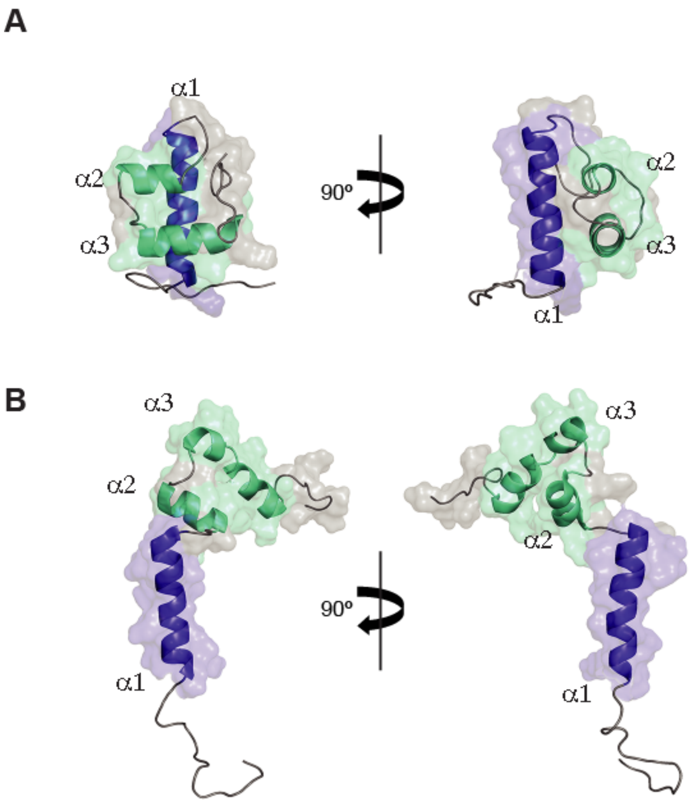


**Fig. S2. Our ComGC$_{SP}$ structure differs markedly from a previously published one (PDB 5NCA).** Cartoon representations of the two averaged solution structures, face and side views, are shown. We used the same nomenclature and colour scheme than in Fig. 4 and Fig. 5. (**A**) Our ComGC$_{SP}$ structure. (**B**) PDB 5NCA ComGC$_{SP}$ structure.

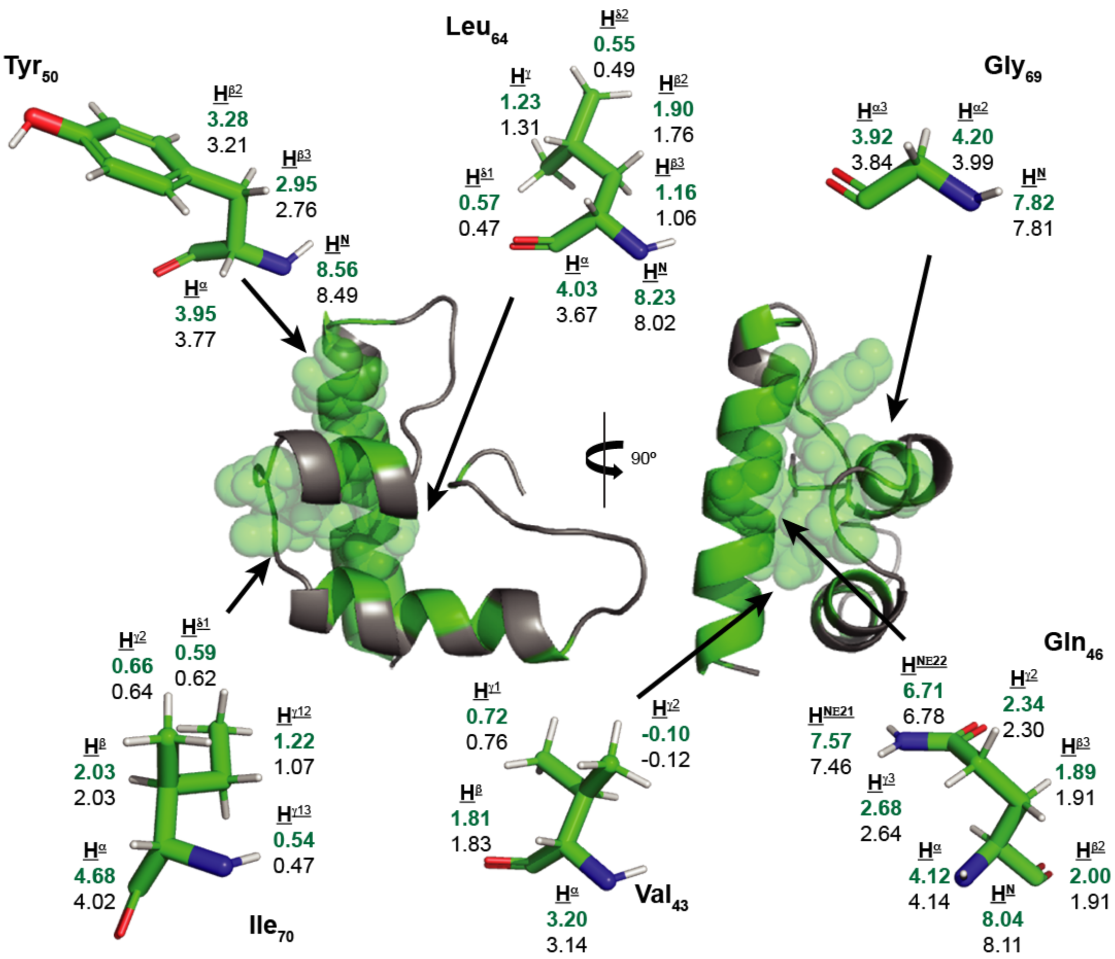


**Fig S3. Conserved residues contributing to the globular fold of ComGC show unusually high chemical shift conservation between ComGC$_{SS}$ and ComGC$_{SP}$ structures.** Cartoon representation of the ordered residues of ComGC$_{SS}$, where conserved residues are highlighted in green, with residues Val$_{43}$, Gln$_{46}$, Tyr$_{50}$, Leu$_{64}$, Gly$_{68}$ and Ile$_{70}$ shown with space filling representation. For these residues, $^{1}$H chemical shifts are shown for ComGC$_{SS}$ (top in bold green) and ComGC$_{SP}$ (bottom).

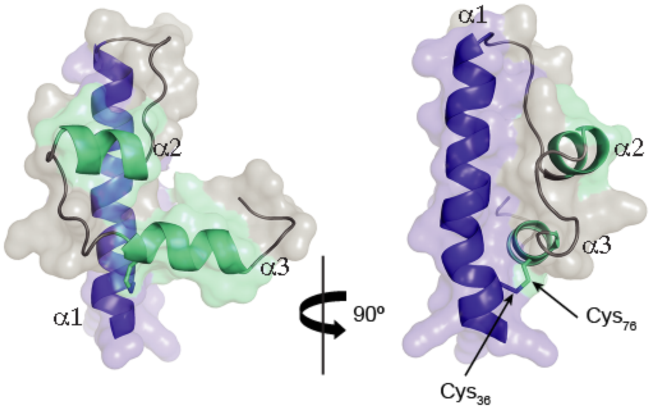


**Fig. S4. 3D model of the globular $ComGC_{BS}$ identifies the disulfide bond known to stabilise that pilin.** $ComGC_{SS}$ has been used as a template to generate a structural model of the globular head of $ComGC_{BS}$. The two cysteines known to form a disulfide bond, $Cys_{36}$ and $Cys_{76}$, are found in close proximity and are expected to significantly stabilise the globular fold.

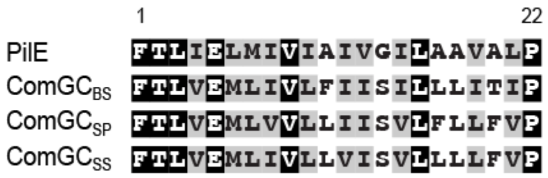


**Fig. S5. The N-terminus in ComGC pilins is highly similar to that of other bacterial T4F major pilins.** Sequence alignment of the N-termini of processed $ComGC_{SS}$, $ComGC_{SP}$, $ComGC_{BS}$, and PilE from *N. gonorrhoeae* T4P. Residues were shaded in black (identical), grey (conserved) or unshaded (different).

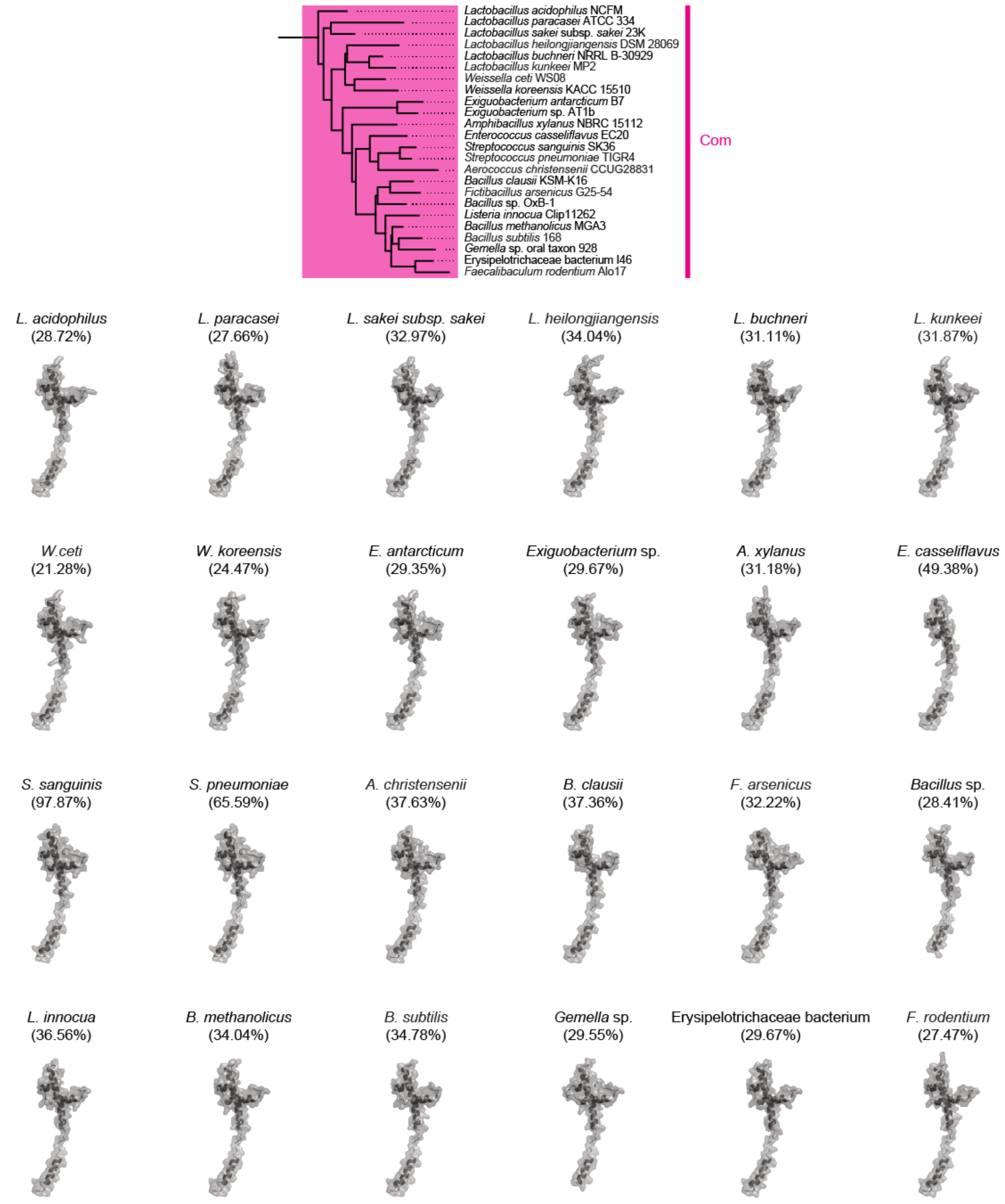


**Fig. S6. Full-length 3D models of ComGC from species used to generate the phylogenetic tree in Fig. 3.** The 3D structures have been modelled using as template (i) our averaged *S sanguinis* ComGC$_{SS}$ structure for the globular head, and (ii) the PulG pilin from *K. oxytoca* for the protruding α1N.

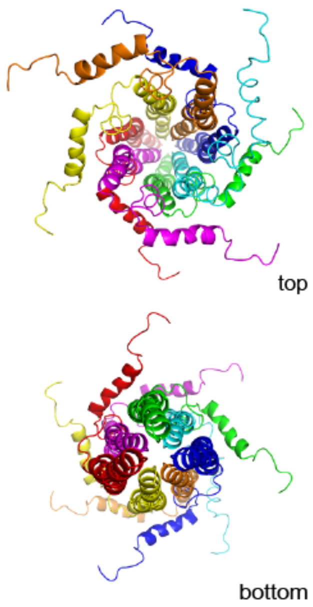


**Fig. S7. 3D model of ComGC pili using the previously published (PDB 5NCA) ComGC$_{SP}$ structure.** Introducing the previous ComGC$_{SP}$ structure into our SWISS-MODEL of ComGC filaments shows a high degree of steric clashes.

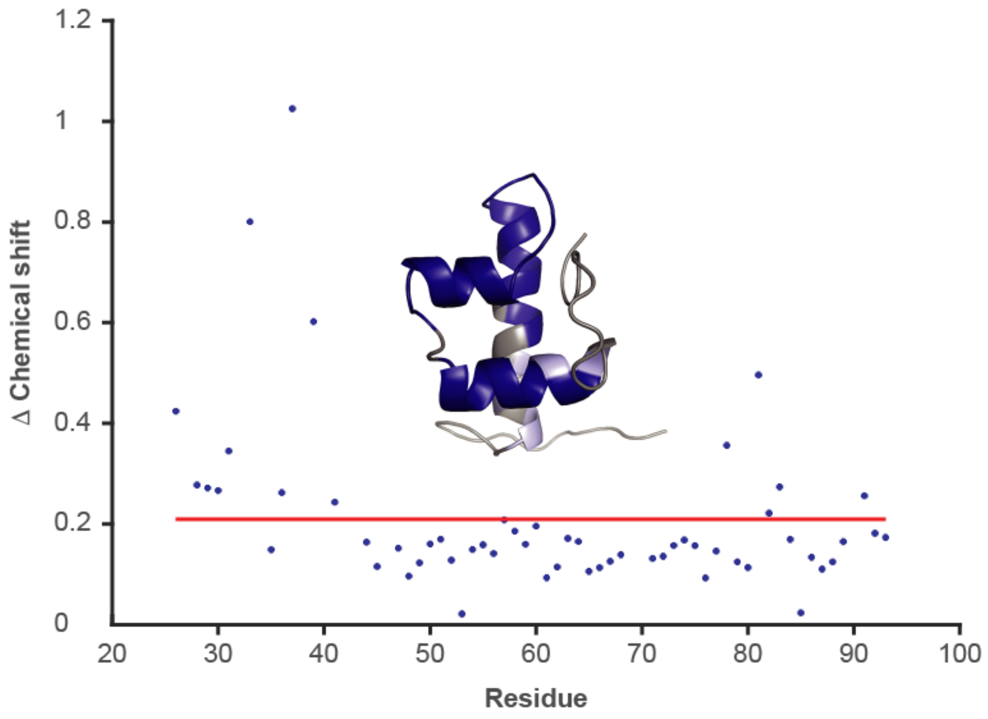


**Fig S8. Amide chemical shift differences between our ComGC$_{SP}$ structure and the previously published one (PDB 5NCA).** Amide chemical shift differences [$\omega_H$+($\gamma_N$/$\gamma_H$)$\omega_N$] are presented against residue number. The average chemical shift difference is indicated as a red line. Those residues for which the difference in chemical shift is greater than the average ± standard deviation are highlighted onto our averaged ComGC$_{SP}$ structure in light blue (inset).

**Supplemental Spreadsheet 1.** List of all the Com systems detected by a search of the RefSeq database (from April 2019) using MacSyFinder with the corresponding Com model.

**Supplemental Spreadsheet 2.** List of all the species that have the potential to express a Com pilus.
